# Stretch-Induced Uncrimping of Equatorial Sclera Collagen Bundles

**DOI:** 10.1101/2022.09.13.507860

**Authors:** Ning-Jiun Jan, Po-Yi Lee, Jacob Wallace, Michael Iasella, Alexandra Gogola, Ian A. Sigal

## Abstract

Stretch-induced collagen uncrimping underlies the nonlinear mechanical behavior of the sclera according to what is often called the process of recruitment. We recently reported experimental measurements of sclera collagen crimp and pressure-induced uncrimping. Our studies, however, were cross-sectional, providing statistical descriptions of crimp with no information on the effects of stretch on specific collagen bundles. Data on bundle-specific uncrimping is necessary to better understand the effects of macroscale input on the collagen microscale and tissue failure. Our goal in this project was to measure bundle-specific stretch-induced collagen uncrimping of sclera. Three goat eyes were cryosectioned sagittally (30μm). Samples of equatorial sclera were isolated, mounted to a custom uniaxial stretcher and imaged with polarized light microscopy at various levels of clamp-to-clamp stretch until failure. At each stretch level, local strain was measured using image tracking techniques. The level of collagen crimping was determined from the bundle waviness, defined as the circular standard deviation of fiber orientation along a bundle. Eye-specific recruitment curves were then computed using eye-specific waviness at maximum stretch before sample failure to define fibers as recruited. Nonlinear mixed effect models were used to determine the associations of waviness to local strain and recruitment to clamp-to-clamp stretch. Waviness decreased exponentially with local strain (p<0.001), whereas bundle recruitment followed a sigmoidal curve with clamp-to-clamp stretch (p<0.001). Individual bundle responses to stretch varied substantially, but recruitment curves were similar across sections and eyes. In conclusion, uniaxial stretch caused measurable bundle-specific uncrimping, with the sigmoidal recruitment pattern characteristic of fiber-reinforced soft tissues.

## 1. Introduction

Collagen is a basic component of the eye, and plays a central role in determining tissue biomechanics [1-4]. Changes in ocular collagen have been associated with natural processes like development and aging and with several eye disorders such as myopia, keratoconus and glaucoma [4-12]. To fully understand these processes and disorders and identify, prevent and/or revert their negative effects it is therefore important to understand collagen, and in particular its biomechanics.

The collagen fibers of the eye [13-17], like those of other soft tissues, have natural undulations or waviness called crimp [18-20]. Crimp is central to the nonlinear biomechanics of soft tissue through a process known as fiber recruitment [21-23]. Experimental crimp measurements from tissues like tendon [24], ligament [25], and arterial tissue [26] are consistent with this explanation.

In the ocular biomechanics community, the importance of crimp and recruitment has also been recognized for several decades. Crimp in the eye has been visualized using several imaging modalities, including electron microscopy [15], second harmonic generated (SHG) imaging [27], and magnetic resonance imaging [13]. For the cornea, the crimp morphology and recruitment have recently been quantified using transmitted electron microscopy and uniaxial tension testing, respectively [28]. However, far less is known about collagen recruitment in the sclera. To fill this gap in information, inverse numerical models have been used to predict the collagen crimp structure and recruitment of the sclera [29-32]. The model predictions from these models, while reasonable, were not confirmed experimentally.

Recently, we reported experimental measures of peripapillary sclera collagen crimp [17, 33] and recruitment [16, 34, 35]. We compared the crimp from different regions of the eye and characterized population crimp trends with changes in intraocular pressure (IOP). From these measurements, we were able to derive recruitment curves for both lamina cribrosa and peripapillary sclera. However, the measurements for these studies were obtained from fixed tissues, and therefore crimp could only be quantified at one stretch state or IOP level. The results were cross-sectional, and thus lacked information on how a specific collagen bundle changes with stretch. A characterization of crimp and recruitment tracking bundle-specific changes in fresh tissue is needed to further the development of accurate fiber-based constitutive models of the eye and to understand the effects of macroscale input on the collagen microscale and tissue failure. Our goal in this study was to use unfixed scleral tissue to characterize the stretch-induced collagen bundle uncrimping and recruitment. Specifically, we analyzed the changes in collagen bundle crimp in unfixed equatorial sclera of goat eyes subjected to uniaxial stretch.

## 2. Methods

We prepared longitudinal unfixed sections of equatorial sclera and imaged them using polarized light microscopy (PLM) while the tissue was stretched to failure. The PLM images were used to quantify collagen orientation and from the deformations quantify collagen waviness. Analyzing the PLM images using digital image correlation we also tracked tissue deformations, to compute local strain and to measure clamp-to-clamp stretch. We analyzed the uncrimping of several collagen bundles to test the association between collagen waviness and local strain and to construct recruitment curves under the clamp-to-clamp stretch.

It is important to note that collagen architecture is hierarchical and complex [4]. Even in tendon, where the collagen is much more clearly organized compared to the eye, there is sometimes disagreement on the specific meaning of various terms to describe the hierarchy of the collagen architecture [36]. To avoid confusion, we clarify that for the purposes of this paper, we use the term collagen “bundles” to refer to groups of contiguous and aligned collagen fibers or fibrils. We have shown that these groupings are discernable in sclera using PLM [33].

### 2.1 Sample preparation and mounting

Three fresh goat (caprine) eyes were acquired from a local abattoir and processed within 8 hours of death following previously described methods [16, 17, 37, 38], with some modifications for longitudinal sectioning of unfixed whole globes rather than coronal sections of fixed optic nerve heads. Briefly, using scalpels, forceps, and razors, extraneous muscle, fat, and episcleral tissue were removed. Each eye was mounted in an embedding cryomold and filled with optimal cutting temperature compound. All eyes were aligned within the molds in the nasal-temporal and superior-inferior anatomical directions for sectioning. After embedding, the eyes were flash frozen in isopentane chilled in liquid nitrogen (−176°C). The molds were stored in plastic bags at -80° C until cryosectioned sagittally into 30 μm sections. Using a standard anti-roll plate and cold fine-tip brush, sections were held flat before transferring to an uncharged glass slide. The section was washed with PBS three times to remove the cryoprotectant agents. Over 20 consecutive sections through the ONH and cornea were collected for each eye. Of those, two consecutive sections free of artifacts, such as folds or breaks, were used for analysis. Overall, 6 sections from three goat eyes were tested. Centimeter-long strips of equatorial sclera were carefully isolated with a razor blade and transferred to a modified uniaxial stretcher (Microvice, S.T. Japan, FL, USA) (**Fig. 1A**). The tissue section was mounted by clamping the anterior and posterior ends (**Fig. 1B**). The section was again twice washed with PBS.

**Figure 1.**
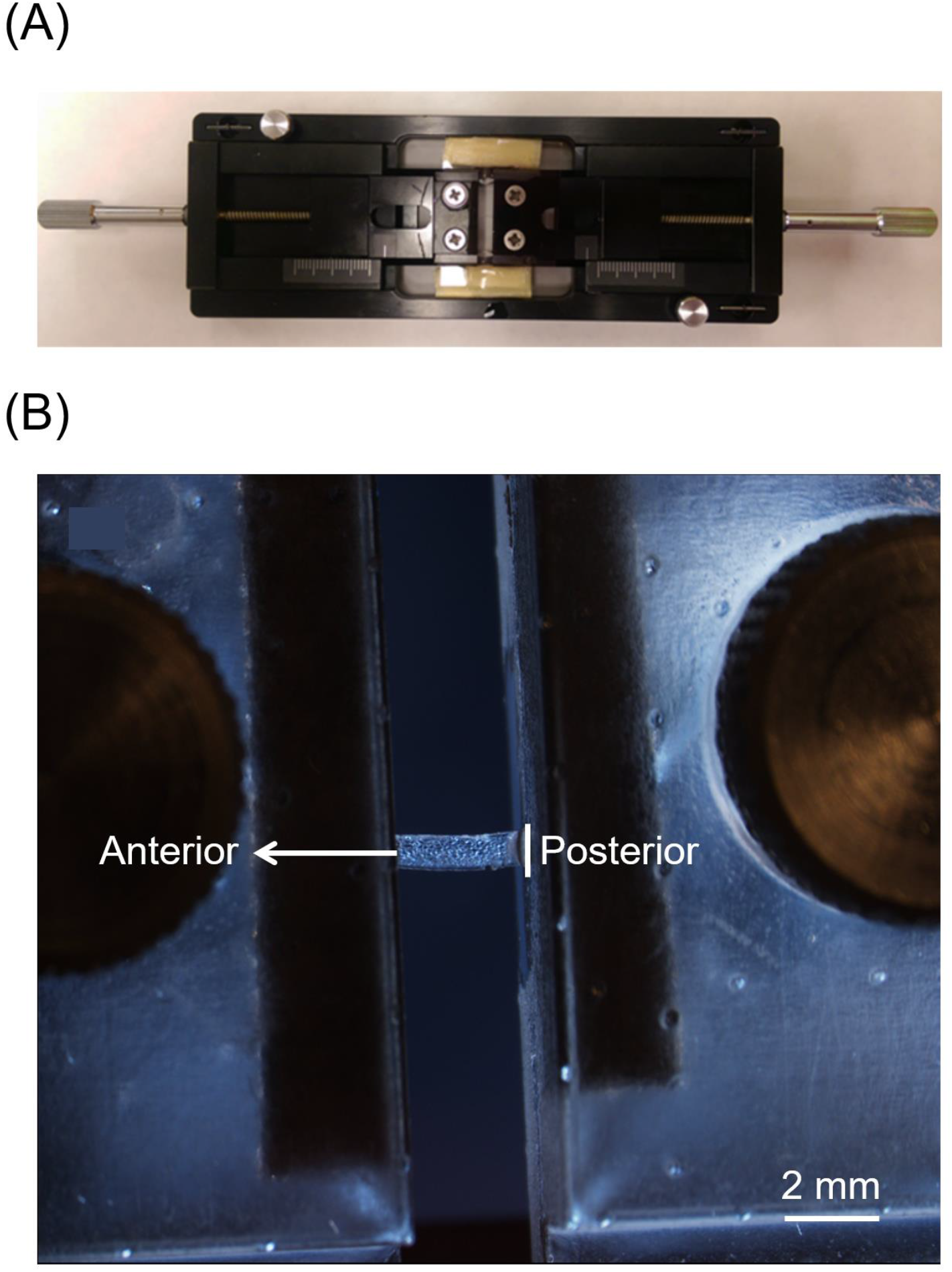
A modified uniaxial stretcher was used to test unfixed equatorial scleral sections. (A) Photograph of the device. The modifications made to keep the sample hydrated are discernible above and below the clamps. (B) Close-up of the clamped tissue from the underside with the custom clamps. The clamp holding the posterior side was fixed while the clamp holding the anterior side was translated to induce stretch. The large thumb-screws used to tighten the clamp are visible on the sides. The photo was acquired at a slightly tilted angle to facilitate discerning the clamping side on the posterior side. This occluded the clamping side on the anterior side. Tests were carried out with an orientation that allowed visualization of both clamp sides to measure clamp-to-clamp distance.

### 2.2 Imaging

The mounted sections were imaged at various stretch levels using PLM as previously described [16, 17, 37, 38], with some modifications for imaging unfixed mounted sections (**Figs. 2, 3, and 4A**). Briefly, a polarizer filter was placed above the sample and an analyzer filter below the sample (Hoya, Tokyo, Japan). An Olympus SZX16 microscope (6.3x magnification setting) paired with an Olympus DP80 camera (36-bit, RGB, pixel-shift setting) was used to acquire the images. (Olympus, Tokyo, Japan) Images had a pixel size of 0.68 μm using a 0.8x objective (numerical aperture, 0.12). Four images were acquired, each with filter orientations 45° apart from each other. Local collagen fiber orientation was determined using the relative changes in intensity at each pixel (**Fig. 3A**) [37, 39]. For all visualizations, the pixel intensities were scaled using an energy, a parameter previously described [37]. Energy is helpful to distinguish regions with collagen from regions outside the sample. This, in turn, was helpful to discern tissue texture. High energy values also indicate that the fibers are in the plane of the section [40]. This was extremely important because the mechanical response of fibers perpendicular to the section is likely more complex and not the target of our study.

**Figure 2.**
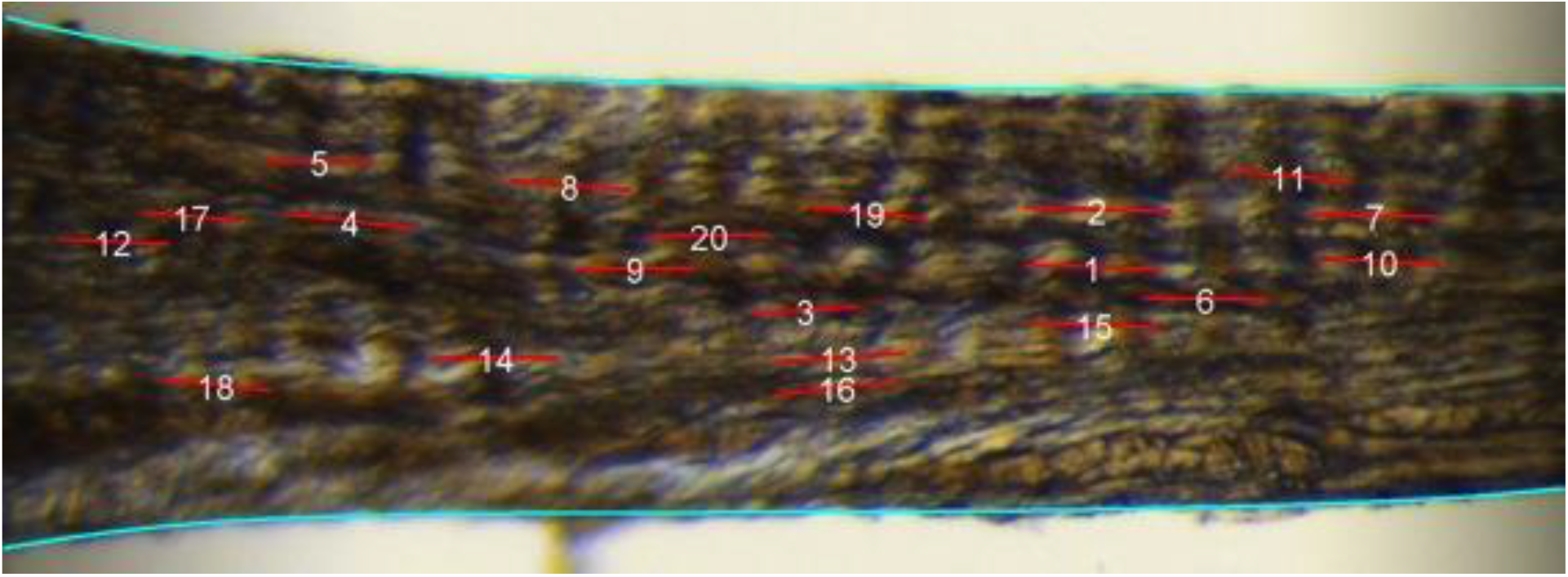
Example PLM image. Collagen bundles run from left to right in the image. Vertical light and dark bands indicative of collagen waviness are clearly discernible. Collagen bundles were marked manually (red lines, numbered) to track the bundle-specific waviness and local strain. Waviness was calculated using the circular standard deviation of the collagen fiber orientations along each line markers.

**Figure 3.**
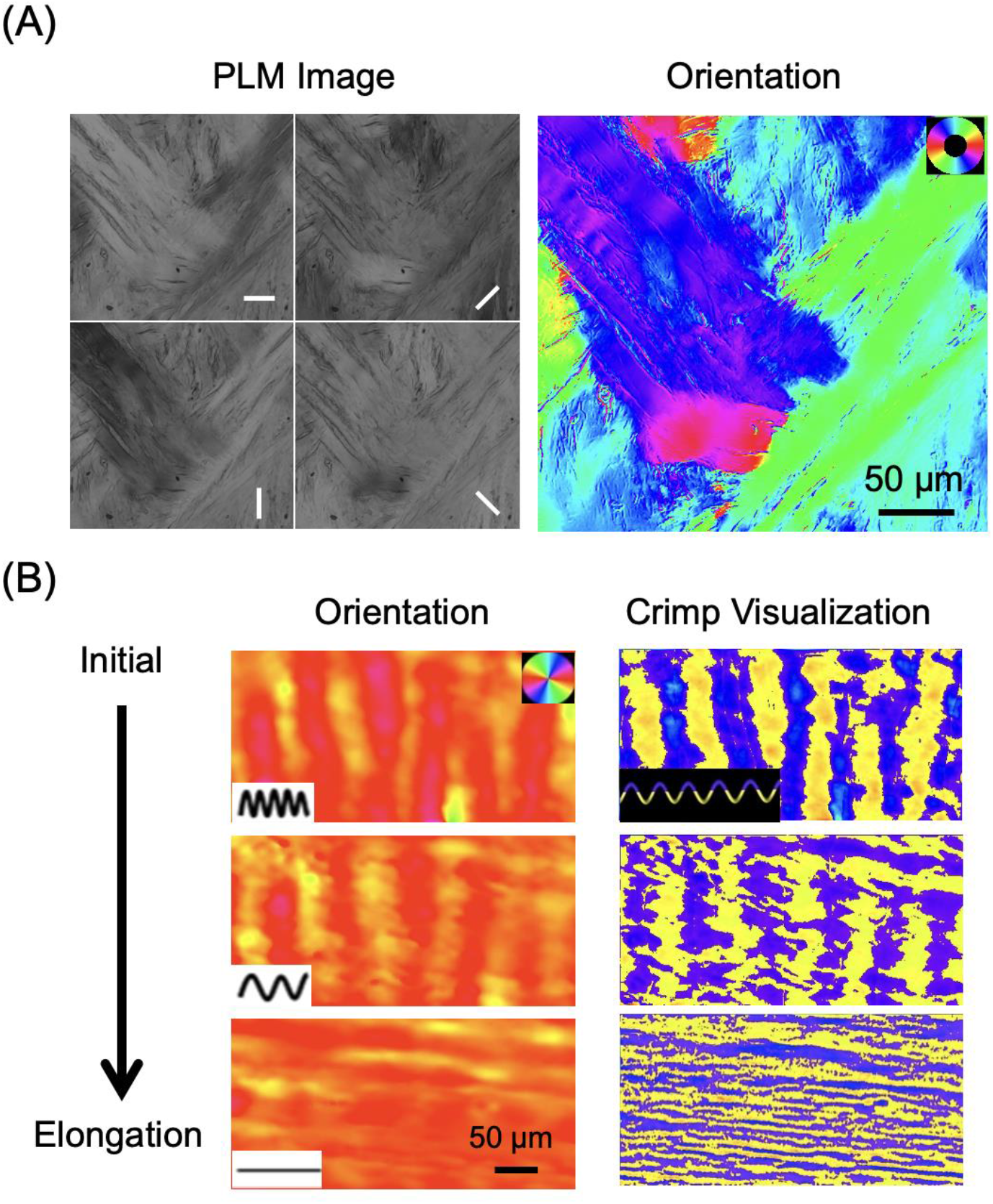
Collagen orientation calculation and crimp visualization from PLM images. (A) Four PLM images were used to determine the local collagen fiber orientation at each pixel. White bars represent the polarization states. (B) Orientation plots (left) and crimp visualization (right). The orientation plots directly color each pixel with the local orientation. The color bands indicate the undulating nature of the fibers, in this case running horizontally. If the fibers were straight, there would be lines of uniform color. To help visualize the crimp we processed the orientations as described elsewhere. This processing results in yellow/purple bands perpendicular to the fiber direction. These correspond to the crimp. Three stages of uncrimping are shown from top to bottom. As the stretch increased, the undulation decreased, suggesting the gradual uncrimping of the collagen bundles.

**Figure 4.**
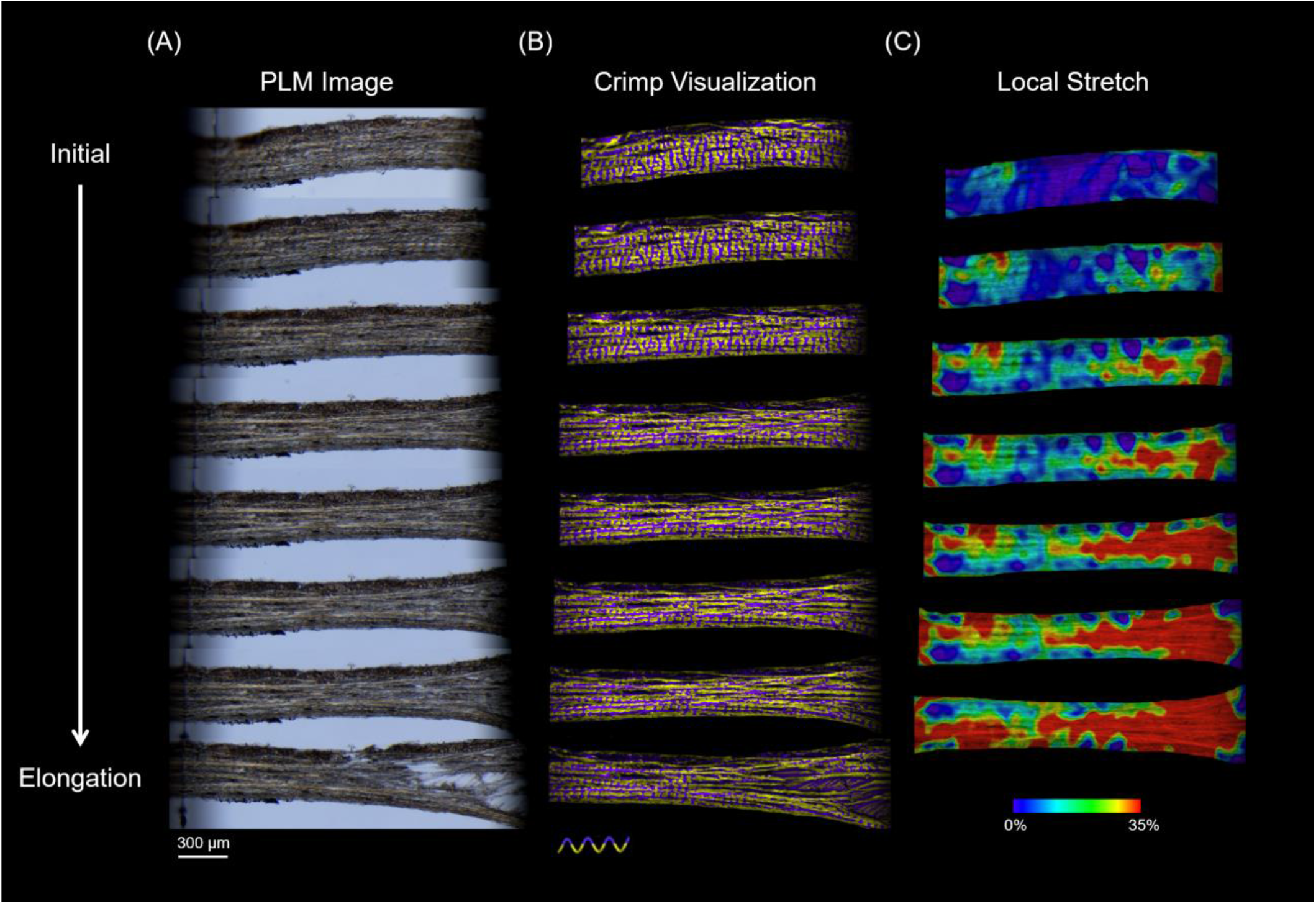
Example image sequence of equatorial sclera at various levels of uniaxial stretch, increasing from initial at the top to failure at the bottom. (A) Raw PLM images. Four PLM images at different polarization states were used to determine local collagen fiber orientation at every pixel. (B) The orientation maps were processed as described in the main text into yellow/purple bands to simplify visualization of collagen crimp. (C) The images were analyzed using tracking routines to calculate the local strain. Note that the strain maps were computed between two consecutive stretch states, and therefore there was one fewer image of strain that raw images or crimp. The overall increase in sample length is discernible, as is the quick loss of initial curvature. At low stretch the yellow-purple bands perpendicular to the sample main orientation are clear indications of crimp. Stretch resulted in the loss of crimp, on some regions first, then in others. Even at the time of maximum stretch (right before failure, second row from the bottom), there were still bundles that still exhibited some crimp. Note that these bands are discernible even for very small crimp and it is likely that the crimp in these regions is already much smaller than before sample stretch.

### 2.3 Collagen waviness

From the PLM-derived orientations, we measured the waviness of collagen fiber bundles. We quantified the waviness using the circular standard deviation [41] of the collagen fiber orientations along a collagen bundle, as described previously [16]. Briefly, a straight fiber bundle would have a constant angle value, and therefore the waviness would be 0°. On the other hand, a wavy fiber bundle would have variable angle values throughout the fiber bundle, and therefore the waviness would be greater than 0°. To measure the waviness, we sampled the orientations using line segments placed along the length of a collagen bundle. This same line segments were used to measure the bundle waviness and strain (**Fig. 2**). Analysis was only performed on bundles that were identifiable throughout all stretch levels. To visualize the crimp changes with stretch, we used a previously described algorithm for highlighting the crimp (**Figs. 3B and 4B**) [16]. Briefly, pixels were colored purple or yellow depending on whether the orientation was larger or smaller than the local average orientation, respectively. In crimped collagen bundles, the result was clear bands of alternating purple and yellow, each corresponding to half a crimp period. Uncrimped, or recruited, bundles show no be clear bands. It is important to note that we have shown that this technique for analyzing collagen fiber crimp does not require discerning fiber undulations or even the edges of the fibers or bundles [10]. The PLM-derived orientation resolved at a pixel encodes sub-pixel information on fiber orientation. Thus, it is possible to use a straight line along a bundle to derive highly accurate measurements of bundle undulations without the need to distinguish visually the bundle edges or the undulations. This is extremely important at high levels of stretch where edges and undulations become essentially impossible to distinguish, yet the signal remains adequate.

### 2.4 Tissue stretching

The clamp holding the posterior side was fixed while the clamp holding the anterior side was translated to cause stretch (**Fig. 1B**). The clamps were constrained to displace in the anterior-posterior direction, along the main axis of the sample. Each level of stretch was held for 15 min to allow vibrations or viscoelastic effects to dissipate before imaging. Each sample was stretched until a visual rip in the tissue was observed. Throughout the imaging session the sample was kept hydrated with 1X neutral phosphate buffered saline. Each section was stretched between five and seven times, including the last one where they failed. The average total clamp displacement at failure was 462 μm and the average step size was 82 μm.

### 2.5 Clamp-to-clamp stretch

The clamp-to-clamp stretch was defined as percent change in clamp-to-clamp distance as the sample was stretched, and represented how much the entire tissue strip was stretched. The distance was measured by segmenting the border between the clamp and the tissue in each image and determining the shortest distance between the two edges for each stretch state. The percent change in distance was calculated relative to the distance between the clamps in the last stretch state before failure. With this definition the clamp-to-clamp stretch was 0% before testing, and 100% at maximum stretch before failure.

### 2.6 Local strain

The local strain was a measure of how the tissue locally deformed as a result of the uniaxial stretching. This strain was measured by how much the tissue locally stretched in the direction of the largest stretch, also known as the first principal strain. The strains were calculated using an image registration algorithm previously reported [42]. Briefly, the “before” image is morphed to register it with the “after” image. The strains are then calculated based on how each pixel was morphed to get to the final image. The strains were calculated as a percentage, where 0% would indicate no change in length and 100% would indicate doubling in length. We used the first principal strain as it represents the magnitude of the largest stretch. The local strain of a specific bundle was defined as the average strain over a line segment manually drawn along the bundle. To visualize the local strain, we pseudo-colored the raw images using a rainbow color scheme, where small strains are colored violet and large strains red and the pixel intensities were weighted using an energy at the corresponding pixel (**Fig. 4C**).

### 2.7 Recruitment

Recruitment curves were computed based on the changes in collagen waviness, in a similar way as in our previous report, [16], with a few adjustments for tracking bundle-specific changes. Briefly, we tracked the waviness in, at least, eighteen collagen fiber bundles in each section through at least four levels of stretch before failure. The bundles were selected to be spread over the sample. The percentages of recruited collagen bundles were calculated for each level of clamp-to-clamp stretch for each tissue section. We defined eye-specific waviness thresholds to determine whether a collagen fiber bundle was considered recruited. We set this threshold as the 75^th^ percentile of the waviness at the final stretch point for each eye. A recruitment curve was constructed for each section of each eye. In our previous study on recruitment we observed that changes on this threshold affect the fraction of fibers or bundles considered recruited, but only had a minor effect on the shape of the recruitment curves [16]. Our interpretation of the finding in that study suggests that the threshold is smaller than 100% because the fibers or bundles exhibit a natural curvature distinct from crimp that is also detected by the waviness analysis. This is consistent with the anterior-posterior collagen fibers in the sclera that likely have some curvature associated with the overall globe shape.

### 2.8 Statistical analyses

All statistical analyses were done using the NLME package in R [43].

#### Waviness association with local strain

A mixed effect exponential model accounting for autocorrelations of measurements from the same section and eye were used to determine whether, for each collagen bundle, waviness was related to local strain. Mixed effect models incorporate fixed variables (waviness and local strain) with random variables that may affect the sampling population (section and eye) [44].

#### Recruitment curve fitting

Using a mixed effect model, a sigmoidal curve was fit to the measurements from all the sections of percent of bundles recruited as a function of percent total clamp-to-clamp stretch. The mixed effects model accounts for autocorrelations of measurements from the section and eye.

## 3. Results

A total of 97 collagen bundles were tracked across all six sections of the three eyes. At baseline (mounted, before stretching), the average bundle wavinesses were 8.51° for Eye 1, 7.24° for Eye 2, and 5.69° for Eye 3. At the largest stretch before failure, the average bundle wavinesses had decreased to 2.37° for Eye 1, 2.88° for Eye 2, and 2.37° for Eye 3. The waviness versus local strain is shown in **Fig. 5**. For all eyes, waviness exponentially decreased with local strain (P<0.001).

**Figure 5.**
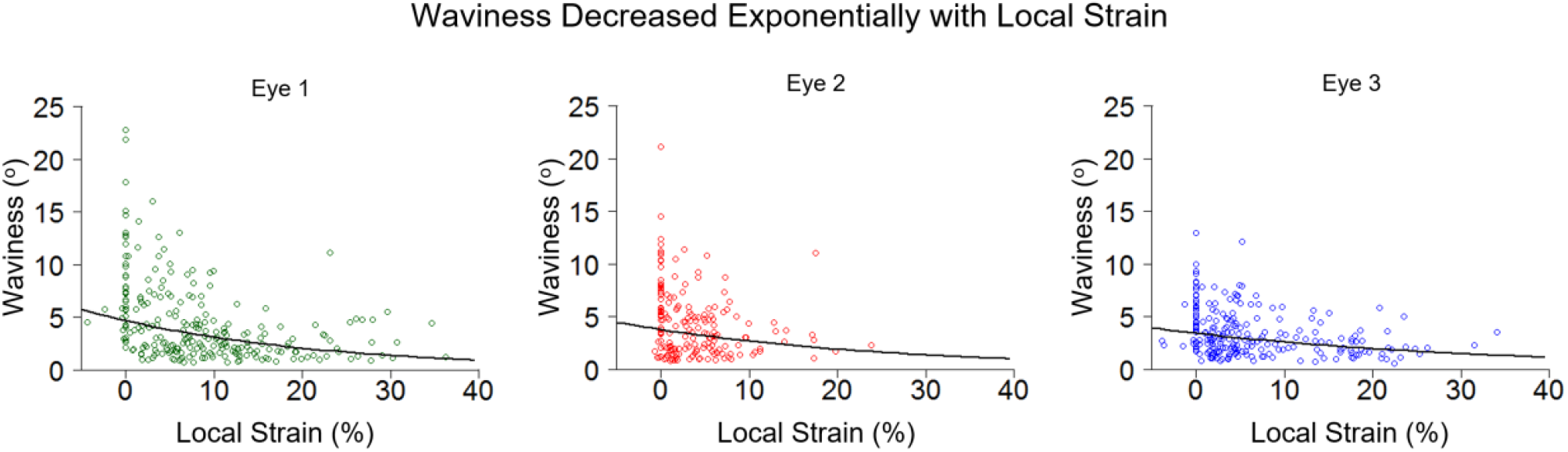
Waviness versus strain in each eye. Each point indicates a bundle measurement of waviness at a given local strain. All three eyes had similar trends, where the waviness decreased exponentially with local strain. Best fit exponential lines are shown in black. All three were significant (P<0.001). At the initial state, with strain zero, every bundle had an initial waviness. Through the test the bundles undergo stretching that changes their waviness. This is the reason for the column of points at zero strain. In a few locations the complex structure of the sclera produced negative local strain (a compression).

The recruitment curves were similar across sections and eyes. Most sections had 10-20% bundles recruited at the base condition, and 70-90% at the final stretch state before failure. The percentage of recruited bundles with clamp-to-clamp stretch followed the sigmoidal pattern expected for fiber-based soft tissues (P<0.001, **Fig. 6**)

**Figure 6.**
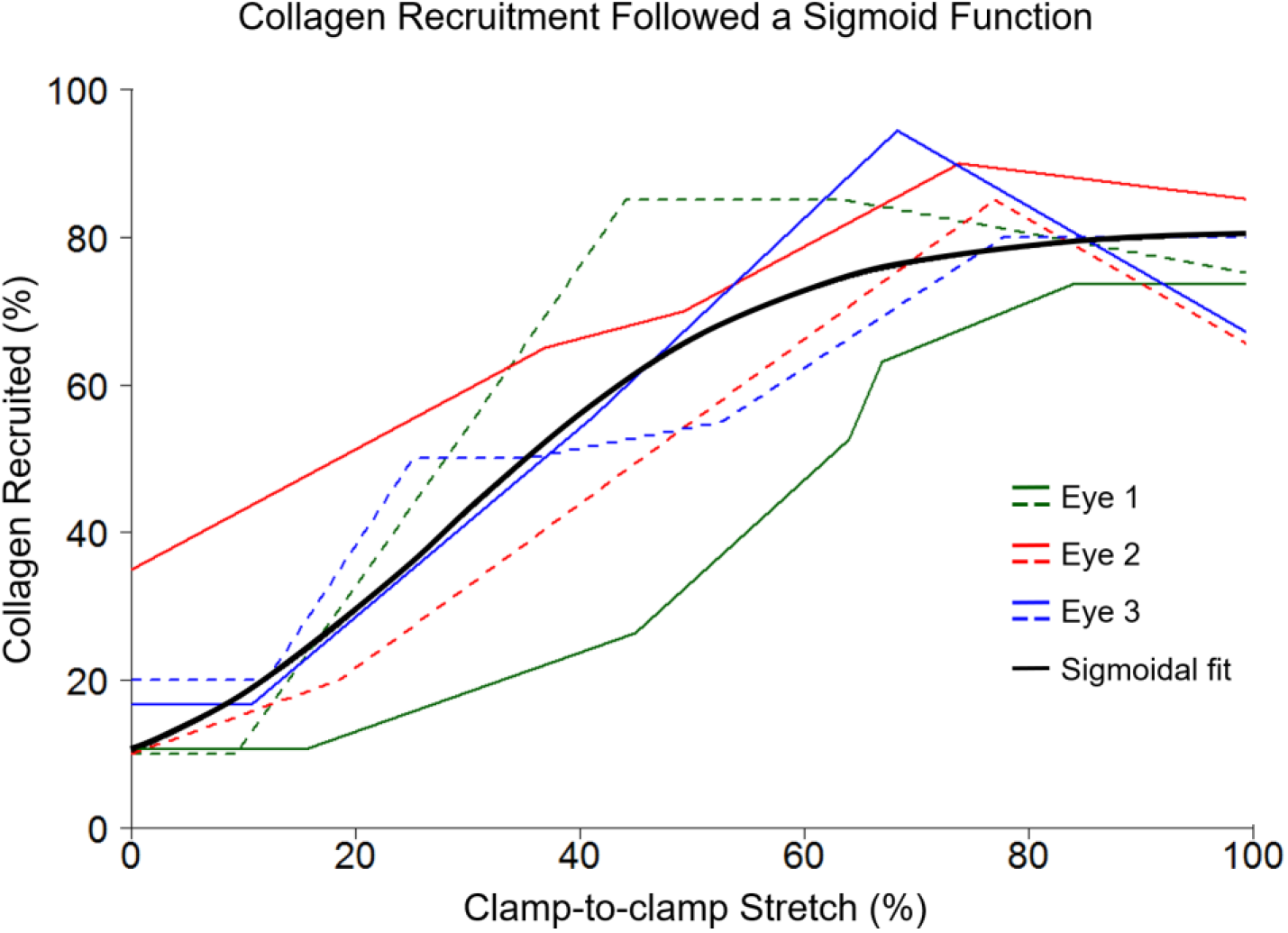
Collagen recruitment curves. The percentage of recruited collagen bundles over the different levels of clamp-to-clamp stretch plotted for two sections (solid and dashed lines) per eye for the three eyes (green, red, and blue). A sigmoidal function was fit across all the measurements (black line). Some sections recruited at lower levels of stretch (shifted to the left) or recruited faster (steeper curves), but overall they all exhibited clear evidence of recruitment.

## 4. Discussion

Our goal was to characterize the stretch-induced collagen bundle uncrimping and recruitment in unfixed goat equatorial sclera subjected to uniaxial stretch. Two main findings arise from our study: i) collagen bundle waviness decreased with increasing local strain; and ii) collagen bundle recruitment followed a sigmoidal pattern. Let us discuss each of these in turn.

### Collagen waviness decreased with local strain

We found that the waviness of collagen bundles decreased as the strains of the corresponding bundles increased. This means that as the crimped fiber bundles were stretched, the bundles became less wavy. This is consistent with the idea that there exists a direct relationship between stretch and bundle uncrimping [1, 3, 16, 23]. By tracking individual collagen bundles with stretch, we experimentally showed this direct relationship. To the best of our knowledge this is the first time that individual sclera collagen bundles have been tracked to measure changes in collagen waviness with stretch. The functional relationship between strain and recruited fibers is critical to the constitutive formulations of sclera biomechanics, and is the microstructural underpinning of the macro-scale nonlinear mechanical properties of the tissue [1, 3, 16, 23]. In addition, we show that the decrease in waviness with stretch follows an exponential curve, whereby the decreases in waviness are progressively smaller as the strain increases. Though this relationship is well-characterized for tendons, ligaments, cornea and other tissues [2, 24-26, 28, 45], it remains still not fully characterized on sclera. Our results herein are consistent with our previous cross-sectional study of collagen waviness in ONHs fixed at various IOPs [16] and with those of other groups [46, 47].

### Collagen bundle recruitment followed a sigmoidal pattern

The collagen bundles recruited over a relatively wide range of clamp-to-clamp stretch, starting to recruit around 10% (toe-region), the rate of recruitment increased to around 50%, and leveling off at around 80% of the final tissue length before failure. The collagen bundles of the equatorial sclera were found to recruit in a sigmoidal function under clamp-to-clamp stretch. The curve of collagen bundle recruitment is closely associated with the change in tissue stiffness [1]. When a sample with homogeneous crimps experiences homogenous strains, the recruitment curve approximates a steep sigmoid function. A steep sigmoid appears more like a step function, resulting in a sudden step change in stiffness of the tissues (**Fig. 7A**). We found, instead, a smooth function over a wide range of strains, indicating progressive recruitment and a smooth nonlinear stiffening over the range of strains in the sclera (**Fig. 7B**). The gradual recruitment results are also consistent with our previous cross-sectional study comparing the crimp of ONHs fixed at various IOPs [16]. Note that the nonlinear response may be the result not only of the variable crimp but of the variable local stretch to which collagen bundles are subjected.

**Figure 7.**
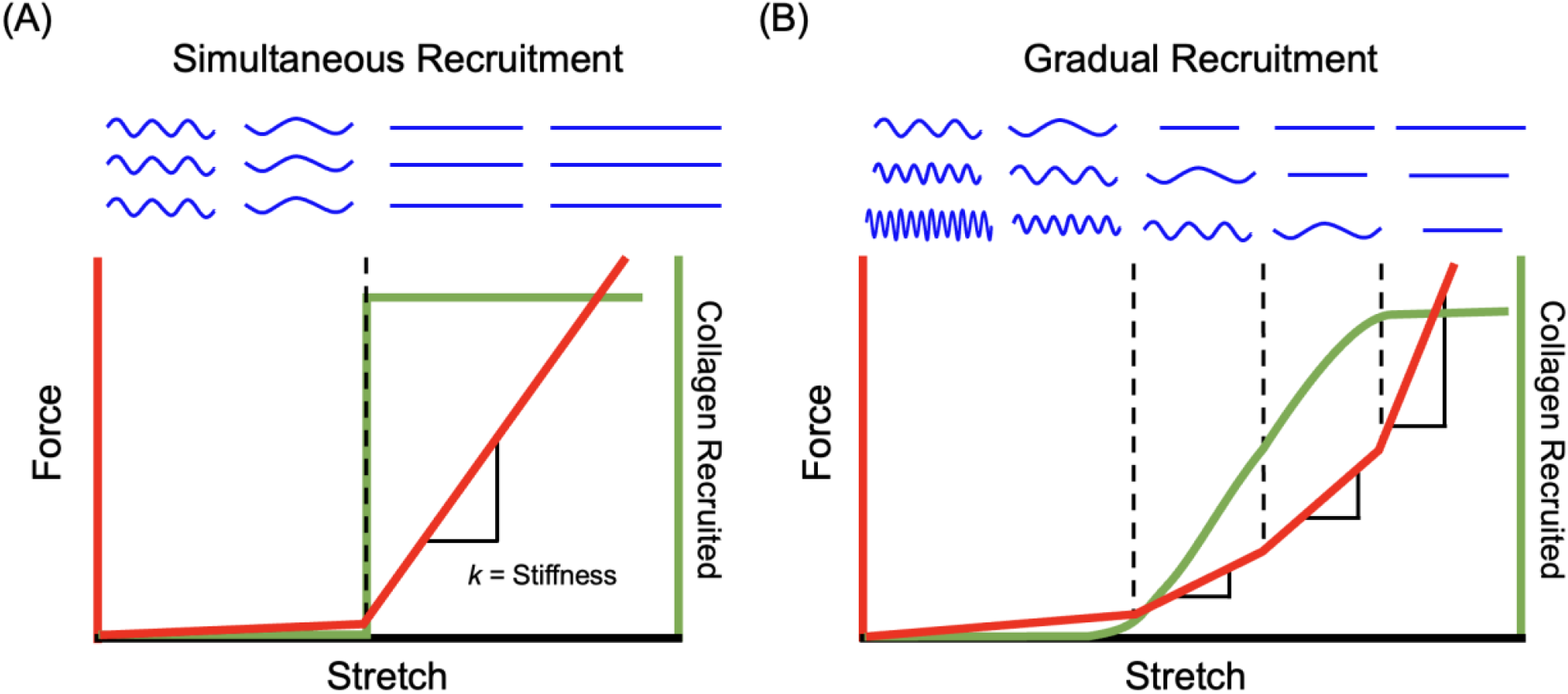
Schematic illustration of step (left) and gradual (right) recruitment of collagen bundles. (A) When all collagen bundles exhibit uniform crimp and stretch the same, the fibers become recruited simultaneously (green curve). Simultaneous recruitment results in an abrupt, step, increase in stiffness (red curve). (B) When collagen bundles exhibit variable crimp or stretch differently, a progressively increasing fraction of fibers become recruited as stretch increases (green curve). Gradual recruitment leads to a nonlinear increase in stiffness (red curve). Figure modified from [17].

It is important to note that our results also suggest that the nonlinear response of the sclera is not only the result of a range of crimp in the collagen bundles. The bundles are also subjected to a range of stretch. Thus, the overall behavior of the tissue is a complex integration of the range of initial crimp and the local stretch to which each bundle is subjected. We do not know all the factors that contribute to determine the local stretch to each bundle, but it seems reasonable to argue that this will likely depend on the complex microarchitecture of the tissue, and therefore that it will be an extremely difficult task. In addition, we must acknowledge that collagen uncrimping and stretch are not the only determinants of sclera biomechanical properties. Other potentially important factors include fiber architecture, density, anisotropy, and other components of the ground matrix, including elastin, glycosaminoglycan and proteoglycan content [48-50], and even hydration [1, 2, 11, 31, 51-54]. These factors may contribute to the variability in how much the waviness changed relative to the amount of local strain experienced by any specific bundle, and to the stretch that each bundle was subjected to. It is possible for example, that some of these bundles first needed to be realigned before they could recruit, while others may have already experienced micro-tears, and the fiber bundles slid past the others, and therefore the crimp remained unchanged. Some bundles lost their waviness and uncrimped at low local strains, whereas others experienced much larger strains before uncrimping. Some collagen bundles never fully uncrimped, even at tissue failure. It is also possible that out-of-plane fibers in a thin tissue section did not bear the load and thus would not recruit under uniaxial stretch. The specific stretch suffered by the collagen bundles may also have been affected by having tested a thin section in which likely most of the fibers had been cut. The distribution of deformations and transfer of forces are potentially quite different in whole tissues in which the fibers form long bundles than when they have been cut [55].

In this study, we quantified the waviness of the collagen crimp, which is but one crimp characteristic. Other crimp characteristics may also be important when characterizing crimp changes with stretch. For example, we have quantified collagen crimp periods previously in ONH tissues as well as around different regions of the eye globe using a simple intuitive measurement method from PLM images [17, 33]. However, as crimped fibers are stretched, the period becomes increasingly difficult to detect until the crimp disappears all together when the fibers become straight. Besides period, other crimp characteristics include amplitude, tortuosity, and maximum deviation angle. We have also previously quantified the crimp amplitude and tortuosity in different regions around the eye globe [33]. Using inverse modeling, the maximum deviation angle has also been predicted for the sclera [29]. Each parameter could have different relationships with stretch. Future studies should consider other crimp characteristics as well as collagen waviness when studying how the sclera responds to load.

Our stretch testing was based on unfixed goat eye tissues without fixation or dehydration, which avoided potentially changing the mechanical properties of tissues and thus affecting the measurements. Typically, in paraffin or plastic embedding methods, the tissue needs to be fixed and dehydrated, which shrinks and warps tissue. In addition, because of the minimal tissue processing, we were able to stretch thin sections of unfixed tissue to observe the crimp changes using PLM. Other studies have used imaging modalities like optical coherence tomography [56], magnetic resonance imaging [57], synchrotron radiation [58] or Raman microspectroscopy [46] to track changes in the eye with IOP. However, these studies have not tracked bundle-specific changes with stretch.

It is important to consider the limitations of our study as well as its strengths. We used goat eyes in this study, which are similar to human eyes in general eye size and shape [59]. However, there are also distinct differences. For example, goat eyes have a thicker sclera than human eyes [60]. It is possible that human tissue would have different crimp morphology and recruit differently than goat tissue, though goat is still important to understand as an animal model [61]. Future work should include other animal models as well as human eyes.

Another limitation is that all PLM orientation analyses were done in 2D sections of the equatorial sclera. It is possible that there is some variability in the sectioning direction, which could affect the consistency of crimp measurements between sections. Also, it is possible some of the collagen had out-of-plane components. To address this, future studies should incorporate 3D collagen fiber orientation measurements using advanced PLM imaging methods [62, 63]. In addition, we did not differentiate between superior and inferior equatorial sclera. Future studies should consider comparing differences between different regions of the eye, like comparing nasal, temporal, superior, and inferior sides of the eye.

It is possible that the tissue processing before testing also affected its mechanical behavior. In particular, the use of cryoprotectants. To reduce this possibility, we washed the tissues multiple times with PBS before testing. The main cryoprotectant agent was Tissue-tek O.C.T. (optimal cutting temperature) compound (Sakura Finetek, California, USA), which has a thick gel-like consistency at room temperature. Since the tests were conducted at room temperature in quasi-static conditions, we posit that the effects of any remaining O.C.T. were small, but they cannot be fully discounted.

Though we have provided a thorough characterization of bundle-specific crimp changes in the sclera with uniaxial stretch, uniaxial stretch testing is a simplified simulation of the tension in the eye induced by IOP. Our current setup using the commercial uniaxial stretcher did not allow us to simulate the more complex conditions in the eye. IOP inflation testing would surely be a much more physiologically realistic situation. However, inflation requires thicker tissues on which it remains a challenge to measure crimp. Structured polarized light microscopy may provide the sufficient resolution and contrast [63].

In conclusion, this study is the first to track bundle-specific collagen crimp changes with stretch in the equatorial sclera. We found that the collagen waviness decreased with local strain and that the collagen bundles recruited in a sigmoidal fashion. Our measurements provide insight into the microstructural basis of nonlinear biomechanics in the sclera. This information helps us develop fiber-based models of the eye, which could help us understand eye biomechanics in relation to aging and diseases like glaucoma.

## Acknowledgments

We would like to acknowledge Ryan O’Malley who helped with histology, Mason Lester who helped with imaging and image processing in this study and Bryn Brazile for helpful discussions. This work was supported, in part, by the National Institutes of Health R01-EY023966, R01-EY028662, T32-EY017271, and P30-EY008098, Research to Prevent Blindness, and the Eye and Ear Foundation (Pittsburgh, Pennsylvania).

## References

[1] Fratzl, P., 2008, Collagen: structure and mechanics, Springer Science & Business Media, New York City, NY.

[2] Holzapfel, G. A., 2001, Handbook of Materials Behavior Models, Academic Press, San Diego, CA.

[3] Grytz, R., and Meschke, G., 2009, “Constitutive modeling of crimped collagen fibrils in soft tissues,” Journal of the mechanical behavior of biomedical materials, 2(5), pp. 522–533.

[4] Boote, C., Sigal, I. A., Grytz, R., Hua, Y., Nguyen, T. D., and Girard, M. J., 2020, “Scleral structure and biomechanics,” Progress in retinal and eye research, 74, p. 100773.

[5] Huang, W., Fan, Q., Wang, W., Zhou, M., Laties, A. M., and Zhang, X., 2013, “Collagen: a potential factor involved in the pathogenesis of glaucoma,” Medical science monitor basic research, 19, p. 237.

[6] Bailey, A. J., 1987, “Structure, function and ageing of the collagens of the eye,” Nature Publishing Group.

[7] Rada, J. A. S., Shelton, S., and Norton, T. T., 2006, “The sclera and myopia,” Experimental eye research, 82(2), pp. 185–200.

[8] McBrien, N. A., Jobling, A. I., and Gentle, A., 2009, “Biomechanics of the sclera in myopia: extracellular and cellular factors,” Optometry and vision science, 86(1), pp. E23–E30.

[9] Campbell, I. C., Coudrillier, B., and Ross Ethier, C., 2014, “Biomechanics of the posterior eye: a critical role in health and disease,” Journal of biomechanical engineering, 136(2), p. 021005.

[10] Gogola, A., Jan, N.-J., Brazile, B., Lam, P., Lathrop, K. L., Chan, K. C., and Sigal, I. A., 2018, “Spatial patterns and age-related changes of the collagen crimp in the human cornea and sclera,” Investigative ophthalmology & visual science, 59(7), pp. 2987–2998.

[11] Coudrillier, B., Pijanka, J., Jefferys, J., Sorensen, T., Quigley, H. A., Boote, C., and Nguyen, T. D., 2015, “Collagen structure and mechanical properties of the human sclera: analysis for the effects of age,” Journal of biomechanical engineering, 137(4), p. 041006.

[12] Albon, J., Karwatowski, W. S., Easty, D. L., Sims, T. J., and Duance, V. C., 2000, “Age related changes in the non-collagenous components of the extracellular matrix of the human lamina cribrosa,” British Journal of Ophthalmology, 84(3), pp. 311–317.

[13] Ho, L. C., Sigal, I. A., Jan, N.-J., Squires, A., Tse, Z., Wu, E. X., Kim, S.-G., Schuman, J. S., and Chan, K. C., 2014, “Magic angle–enhanced MRI of fibrous microstructures in sclera and cornea with and without intraocular pressure loading,” Investigative Ophthalmology & Visual Science, 55(9), pp. 5662–5672.

[14] Newton, R. H., Brown, J. Y., and Meek, K., “Polarized light microscopy technique for quantitatively mapping collagen fibril orientation in cornea,” Proc. BiOS Europe’96, International Society for Optics and Photonics, pp. 278-284.

[15] Andreo, R., and Farrell, R., 1982, “Corneal small-angle light-scattering theory: wavy fibril models,” Journal of the Optical Society of America, 72(11), pp. 1479–1492.

[16] Jan, N.-J., and Sigal, I. A., 2018, “Collagen fiber recruitment: a microstructural basis for the nonlinear response of the posterior pole of the eye to increases in intraocular pressure,” Acta biomaterialia, 72, pp. 295–305.

[17] Jan, N.-J., Gomez, C., Moed, S., Voorhees, A. P., Schuman, J. S., Bilonick, R. A., and Sigal, I. A., 2017, “Microstructural crimp of the lamina cribrosa and peripapillary sclera collagen fibers,” Investigative ophthalmology & visual science, 58(9), pp. 3378–3388.

[18] Franchi, M., Fini, M., Quaranta, M., De Pasquale, V., Raspanti, M., Giavaresi, G., Ottani, V., and Ruggeri, A., 2007, “Crimp morphology in relaxed and stretched rat Achilles tendon,” Journal of anatomy, 210(1), pp. 1–7.

[19] Komatsu, K., Mosekilde, L., Viidik, A., and Chiba, M., 2002, “Polarized light microscopic analyses of collagen fibers in the rat incisor periodontal ligament in relation to areas, regions, and ages,” Anatomical Record, 268(4), pp. 381–387.

[20] Yang, B., Lee, P. Y., Hua, Y., Brazile, B., Waxman, S., Ji, F., Zhu, Z., and Sigal, I. A., 2021, “Instant polarized light microscopy for imaging collagen microarchitecture and dynamics,” Journal of Biophotonics, 14(2), p. e202000326.

[21] Holzapfel, G. A., and Ogden, R. W., 2006, Mechanics of biological tissue, Springer Science & Business Media.

[22] Ethier, C. R., and Simmons, C. A., 2007, Introductory biomechanics: from cells to organisms, Cambridge University Press.

[23] Diamant, J., Keller, A., Baer, E., Litt, M., and Arridge, R., 1972, “Collagen; ultrastructure and its relation to mechanical properties as a function of ageing,” Proceedings of the Royal Society of London. Series B. Biological Sciences, 180(1060), pp. 293–315.

[24] Hansen, K. A., Weiss, J. A., and Barton, J. K., 2001, “Recruitment of Tendon Crimp With Applied Tensile Strain,” Journal of Biomechanical Engineering, 124(1), pp. 72–77.

[25] Thornton, G. M., Shrive, N. G., and Frank, C. B., 2002, “Ligament creep recruits fibres at low stresses and can lead to modulus-reducing fibre damage at higher creep stresses: a study in rabbit medial collateral ligament model,” J. Orth. Res., 20(5), pp. 967–974.

[26] Hill, M. R., Duan, X., Gibson, G. A., Watkins, S., and Robertson, A. M., 2012, “A theoretical and non-destructive experimental approach for direct inclusion of measured collagen orientation and recruitment into mechanical models of the artery wall,” J. Biomech., 45(5), pp. 762–771.

[27] Winkler, M., Chai, D., Kriling, S., Nien, C. J., Brown, D. J., Jester, B., Juhasz, T., and Jester, J. V., 2011, “Nonlinear optical macroscopic assessment of 3-D corneal collagen organization and axial biomechanics,” Investigative Ophthalmology & Visual Science, 52(12), pp. 8818–8827.

[28] Liu, X., Wang, L., Ji, J., Yao, W., Wei, W., Fan, J., Joshi, S., Li, D., and Fan, Y., 2014, “A mechanical model of the cornea considering the crimping morphology of collagen fibrils,” Investigative ophthalmology & visual science, 55(4), pp. 2739–2746.

[29] Grytz, R., and Meschke, G., 2010, “A computational remodeling approach to predict the physiological architecture of the collagen fibril network in corneo-scleral shells,” Biomechanics and Modeling in Mechanobiology, 9(2), pp. 225–235.

[30] Grytz, R., Meschke, G., and Jonas, J. B., 2011, “The collagen fibril architecture in the lamina cribrosa and peripapillary sclera predicted by a computational remodeling approach,” Biomechanics and modeling in mechanobiology, 10(3), pp. 371–382.

[31] Fazio, M. A., Grytz, R., Morris, J. S., Bruno, L., Gardiner, S. K., Girkin, C. A., and Downs, J. C., 2014, “Age-related changes in human peripapillary scleral strain,” Biomechanics and modeling in mechanobiology, 13(3), pp. 551–563.

[32] Girard, M. J., Suh, J.-K. F., Bottlang, M., Burgoyne, C. F., and Downs, J. C., 2009, “Scleral biomechanics in the aging monkey eye,” Investigative ophthalmology & visual science, 50(11), pp. 5226–5237.

[33] Jan, N.-J., Brazile, B. L., Hu, D., Grube, G., Wallace, J., Gogola, A., and Sigal, I. A., 2018, “Crimp around the globe; patterns of collagen crimp across the corneoscleral shell,” Experimental Eye Research.

[34] Lee, P.-Y., Yang, B., Hua, Y., Waxman, S., Zhu, Z., Ji, F., and Sigal, I. A., 2022, “Real-time imaging of optic nerve head collagen microstructure and biomechanics using instant polarized light microscopy,” Experimental Eye Research, 217, p. 108967.

[35] Gogola, A., Jan, N.-J., Lathrop, K. L., and Sigal, I. A., 2018, “Radial and circumferential collagen fibers are a feature of the peripapillary sclera of human, monkey, pig, cow, goat, and sheep,” Investigative ophthalmology & visual science, 59(12), pp. 4763–4774.

[36] Handsfield, G. G., Slane, L. C., and Screen, H. R., 2016, “Nomenclature of the tendon hierarchy: an overview of inconsistent terminology and a proposed size-based naming scheme with terminology for multi-muscle tendons,” J. Biomech., 49(13), pp. 3122–3124.

[37] Jan, N.-J., Grimm, J. L., Tran, H., Lathrop, K. L., Wollstein, G., Bilonick, R. A., Ishikawa, H., Kagemann, L., Schuman, J. S., and Sigal, I. A., 2015, “Polarization microscopy for characterizing fiber orientation of ocular tissues,” Biomedical optics express, 6(12), pp. 4705–4718.

[38] Jan, N.-J., Lathrop, K., and Sigal, I. A., 2017, “Collagen architecture of the posterior pole: high-resolution wide field of view visualization and analysis using polarized light microscopy,” Investigative ophthalmology & visual science, 58(2), pp. 735–744.

[39] Shribak, M., and Oldenbourg, R., 2003, “Techniques for fast and sensitive measurements of two-dimensional birefringence distributions,” Applied Optics, 42(16), pp. 3009–3017.

[40] Yang, B., Jan, N. J., Brazile, B., Voorhees, A., Lathrop, K. L., and Sigal, I. A., 2018, “Polarized light microscopy for 3-dimensional mapping of collagen fiber architecture in ocular tissues,” Journal of biophotonics, 11(8), p. e201700356.

[41] Jammalamadaka, S. R., and Sengupta, A., 2001, Topics in circular statistics, World Scientific Publishing Co.

[42] Sigal, I. A., Grimm, J. L., Jan, N. J., Reid, K., Minckler, D. S., and Brown, D. J., 2014, “Eye-specific IOP-induced displacements and deformations of human lamina cribrosa,” Invest Ophthalmol Vis Sci, 55(1), pp. 1–15.

[43] Team, R. C., 2013, “R: A language and environment for statistical computing. R Foundation for Statistical Computing, Vienna, Austria,” http://www.R-project.org/.

[44] Gałecki, A., and Burzykowski, T., 2013, “Linear mixed-effects model,” Linear Mixed-Effects Models Using R, Springer, pp. 245–273.

[45] York, T., Kahan, L., Lake, S. P., and Gruev, V., 2014, “Real-time high-resolution measurement of collagen alignment in dynamically loaded soft tissue,” Journal of biomedical optics, 19(6), p. 066011.

[46] Chakraborty, N., Wang, M., Solocinski, J., Kim, W., and Argento, A., 2016, “Imaging of scleral collagen deformation using combined confocal raman microspectroscopy and polarized light microscopy techniques,” PLoS One, 11(11), p. e0165520.

[47] Argento, A., Kim, W., Rozsa, F. W., DeBolt, K. L., Zikanova, S., and Richards, J. R., 2014, “Shear behavior of bovine scleral tissue,” Journal of biomechanical engineering, 136(7), p. 071011.

[48] Hatami-Marbini, H., and Pachenari, M., 2020, “Hydration related changes in tensile response of posterior porcine sclera,” Journal of the Mechanical Behavior of Biomedical Materials, 104, p. 103562.

[49] Hatami-Marbini, H., and Pachenari, M., 2021, “Tensile viscoelastic properties of the sclera after glycosaminoglycan depletion,” Current Eye Research, 46(9), pp. 1299–1308.

[50] Hatami-Marbini, H., and Mehr, J. A., 2022, “Modeling and experimental investigation of electromechanical properties of scleral tissue; a CEM model using an anisotropic hyperelastic constitutive relation,” Biomechanics and Modeling in Mechanobiology, pp. 1–13.

[51] Birch, H. L., Thorpe, C. T., and Rumian, A. P., 2013, “Specialisation of extracellular matrix for function in tendons and ligaments,” Muscles, Ligaments and Tendons Journal, 3(1), pp. 12–22.

[52] Ethier, C. R., Johnson, M., and Ruberti, J., 2004, “Ocular biomechanics and biotransport,” Annu. Rev. Biomed. Eng., 6, pp. 249–273.

[53] Rada, J. A., Achen, V. R., Penugonda, S., Schmidt, R. W., and Mount, B. A., 2000, “Proteoglycan composition in the human sclera during growth and aging,” Investigative ophthalmology & visual science, 41(7), pp. 1639–1648.

[54] Gelman, S., Cone, F. E., Pease, M. E., Nguyen, T. D., Myers, K., and Quigley, H. A., 2010, “The presence and distribution of elastin in the posterior and retrobulbar regions of the mouse eye,” Experimental eye research, 90(2), pp. 210–215.

[55] Voorhees, A. P., Jan, N.-J., Hua, Y., Yang, B., and Sigal, I. A., 2018, “Peripapillary sclera architecture revisited: a tangential fiber model and its biomechanical implications,” Acta biomaterialia, 79, pp. 113–122.

[56] Baumann, B., Rauscher, S., Glösmann, M., Götzinger, E., Pircher, M., Fialová, S., Gröger, M., and Hitzenberger, C. K., 2014, “Peripapillary rat sclera investigated in vivo with polarization-sensitive optical coherence tomography,” Investigative ophthalmology & visual science, 55(11), pp. 7686–7696.

[57] Ho, L. C., Sigal, I. A., Jan, N.-J., Jin, T., Wu, E. X., Kim, S.-G., Schuman, J. S., and Chan, K. C., “Microstructural organization and macromolecular contents in fibrous tissues of normal and hypertensive eyes with diffusion tensor imaging and magnetization transfer imaging,” Proc. ISMRM 23rd Annual Meeting & Exhibition cum SMRT 24th Annual Meeting Proceedings.

[58] Coudrillier, B., Geraldes, D. M., Vo, N. T., Atwood, R., Reinhard, C., Campbell, I. C., Raji, Y., Albon, J., Abel, R. L., and Ethier, C. R., 2015, “Phase-contrast micro-computed tomography measurements of the intraocular pressure-induced deformation of the porcine lamina cribrosa,” IEEE transactions on medical imaging, 35(4), pp. 988–999.

[59] Ribeiro, A. P., Santos, N. L., Campos, A. F., Teixeira, I. A. M. d. A., and Laus, J. L., 2010, “Ultrasonographic and ecobiometric findings in the eyes of adult goats,” Ciência Rural, 40(3), pp. 568–573.

[60] Al-Redah, S. A. A., 2016, “Ultrasonographic anatomy of the goat eye,” J. Vet. Med. Sci., 15(1), pp. 160–164.

[61] Chen, Y., Wu, W., Zhang, X., Fan, W., and Shen, L., 2011, “Feasibility study on retinal vascular bypass surgery in isolated arterially perfused caprine eye model,” Eye, 25(11), p. 1499.

[62] Yang, B., Jan, N.-J., Lam, P., Lathrop, K. L., and Sigal, I. A., 2017, “Collagen architecture in the third dimension: 3D polarized light microscopy (3DPLM) for mapping in-plane (IP) and out-of-plane (OOP) collagen fiber architecture,” Investigative Ophthalmology & Visual Science, 58(8), pp. 4825–4825.

[63] Yang, B., Brazile, B., Jan, N.-J., Hua, Y., Wei, J., and Sigal, I. A., 2018, “Structured polarized light microscopy for collagen fiber structure and orientation quantification in thick ocular tissues,” Journal of biomedical optics, 23(10), p. 106001.

